# Enhanced metabolic potentials and functional gene interactions of microbial stress response towards high elevation in freshwater lakes

**DOI:** 10.1101/2020.07.10.198234

**Authors:** Huabing Li, Jin Zeng, Lijuan Ren, Qingyun Yan, Qinglong L. Wu

**Author notes:** Correspondence author: Qinglong L. Wu, or State Key Laboratory of Lake Science and Environment, Nanjing Institute of Geography and Limnology, Chinese Academy of Sciences, East Beijing Road 73, Nanjing 210008, People’s Republic of China.

## Abstract

Elevation has strong influence on microbial community composition, but its influence on aquatic microbial functional genes remains unclear. Here, we compared the functional gene structure of microbial communities in surface water between two low-elevation lakes (LELs, with elevation of ca. 530 meters) and two high-elevation lakes (HELs, with elevation of ca. 4,600 meters) by using a metagenomic approach of Geo Chip-based functional gene arrays. We found significant differences in composition but not in richness of the microbial functional genes between the HELs and the LELs. In the HELs, the microbial communities had higher functional capacities in stress responses than those in LELs, which include cold shock, oxygen limitation, osmotic stress, nitrogen limitation, phosphate limitation, glucose limitation, radiation stress, heat shock, protein stress, and sigma factors genes. We also observed higher metabolic potentials in the degradation of aromatic, chitin, cellulose and hemicellulose in HELs than in LELs. By performing network analyses, we found enhanced interactions and complexity among the co-occurring functional genes in the HELs than those in the LELs in terms of network size, links, connectivity, and clustering coefficients. Notably, more functional genes of stress response played module-hub roles in the network of HELs. Overall, we observed contrasting patterns of microbial metabolic potentials and functional gene interactions in different elevational freshwater lakes, and found that the microbial communities developed functional strategies to cope with the harsh conditions at the high elevational lakes.

**IMPORTANCE:** Elevational patterns of biodiversity have attracted scientific interest in the fields of microbial ecology and biogeography. The influence of elevation on aquatic microbial functional gene structure and their metabolic potentials remains unclear. We compared the functional gene structure of microbial communities in surface water between two low-elevation lakes and two high-elevation lakes with a more than 4,000-meter difference in elevation along a mountainside by using GeoChip 5.0, which covered in total 144,000 gene sequences from 393 functional gene families. We found apparent differences in functional gene structures in lakes between the two different elevations. We also found enhanced metabolic potentials and functional gene interactions for microbial stress response with increasing elevation in freshwater lakes. These results highlighted that limnetic microbial communities could develop functional strategies to cope with harsh conditions towards high elevations.

## INTRODUCTION

Elevational patterns of biodiversity have been attracting scientific interests in microbial ecology and biogeography because of their importance in comprehensively understanding the influences of climate change on ecosystems. A considerable number of studies on altitudinal gradients has focused on terrestrial organism and has revealed that the species richness generally exhibit decreasing or unimodal patterns with the increasing of altitude (e.g., 1, 2). The studies on altitudinal species richness of a few freshwater taxa including aquatic plants (3), phytoplankton (4), rotifers (5), crustaceans (6), stream macro-invertebrates (7) and mollusks (8), have indicated a linear decreasing pattern in species richness with altitude, whereas those of chironomids (9) and stream fish (10) have shown a hump-shaped pattern with altitude. Only a few studies have focused on microbial diversity across elevation gradients in freshwater lakes (11, 12), though microbial communities are the most important groups driving biogeochemical cycles and sustaining the whole system in limnetic systems (13).

Microbial communities play central roles in biogeochemical circulations such as carbon (C), nitrogen (N), and phosphorus (P) cycling (13, 14). With the development of microbial molecular biology technology, microbial phylogenetic and functional diversity patterns along elevational gradients are recently receiving considerable attention in ecology and biogeography, most of which have focused on taxonomy. The limited number of studies on microbial communities living in soils (15-17), on stones in streams (18), on leaf surfaces (17), and in lake waters (11, 12) have indicated that the elevational patterns of microbial species richness varied with elevational ranges and habitat types. Kuang *et al*. (19) has demonstrated that microbial functional metabolic potentials in acid mine drainage were strongly influenced by the changes in microbial species richness and in environmental conditions. However, little is known about the elevational patterns of microbial functional traits. It is crucial to reveal microbial functional traits along elevation gradients, because it can improve our understanding of the influences of climate change on the microbial-related ecosystem processes.

Many of the environmental variables that vary with elevation may affect the functional gene structure of microbial communities. To the best of our knowledge, there is only one study to date has examined the microbial functional gene diversity along an elevation gradient of the grassland in Tibetan (20). Yang *et al* (20) found that, with the increase of elevation from 3,200 m to 3,800 m, soil microbial functional gene structure changed significantly and this change was mostly caused by the variations of environmental variables including decreases in temperature, the concentrations of carbon, and nutrients. The shifts of these environmental factors have been found even more than twice larger in lakes across the elevations from about 530 m to 4,600 (12). Furthermore, differed from grassland, the high-elevation lakes (HELs), especially those lakes with elevations higher than 4,000 m, are characterized by more harsh conditions including high UV, low primary production, more recalcitrant dissolved organic carbon (DOC) than low-elevation lakes (LELs) (12, 21). The harsh conditions in HELs may result in distinct patterns of microbial metabolic potentials compared to those in LELs, which have not been investigated. It has been confirmed that microbial recruitment to exposed niches is strongly on the basis of selection for stress tolerance traits (22), adding evidence to the classical hypothesis of “everything is everywhere, but environment selects” (23). It will be very likely that the microbial communities at the high elevational lakes may have higher functional capacities in stress responses, such as cold shock, radiation stress, and higher metabolic potentials in the degradation of recalcitrant DOC.

In an ecosystem, microorganisms interact with other organisms and their environment to form a complicated network (24), and understanding this complex network over time and space is one of the key issues in ecology (24, 25). Network analysis has been proven a powerful method not only to examine the complex interactions among microbes and identify keystone species in different ecological systems, such as the human gut (26), river (27), marine (28), soils (29, 30), and rhizosphere (31), but also to show the changes of microbial metabolic potentials in the elevated-atmospheric-CO_2_ and oil-contaminated soils (32,33), and the acidic mining lake (34). These network studies provide perspectives on microbial assemblages that cannot easily be found by simple alpha and beta diversities analyses (31). Previous studies found that the network complexity of different microbial functional genes under elevated atmospheric CO_2_ or more stress (e.g., low pH, limited carbon, nitrogen and phosphorus sources) conditions increased in terms of network size, connectivity, and clustering coefficients (32, 34). Until now, however, the patterns of microbial functional gene interactions have not been investigated in lakes with contrasting patterns of environmental variables at different elevations.

In this study, we hypothesized that (1) the microbial functional gene structure (MFS) differs significantly among the lakes with large difference in elevations; (2) the microbial functional potentials of stress response increase with the increasing alpine stress at high elevations; (3) the harsh conditions of HELs would alter the microbial functional molecular ecological network. To test the hypotheses, we focused on four shallow freshwater lakes at comparable elevations, two HELs (at 4,652 m and 4,608 m, respectively) and two LELs (at 530 m and 525 m, respectively), at Siguniang Mountain in Sichuan province, China. We applied a comprehensive functional gene array (GeoChip 5.0) to analyze the functional gene diversity and metabolic potentials of the microbial communities in the four lakes. Random matrix theory (RMT)-based network analysis was performed to compare the interaction patterns of the co-occurring functional genes between the HELs and LELs. Overall, we demonstrated contrasting patterns of freshwater microbial metabolic potentials and functional gene interactions between the HELs and LELs.

## RESULTS

### Microbial functional gene richness

In total, 17,238 functional genes were detected in the 24 samples, of which 14,912 genes were derived from bacteria, 631 genes from archaea, 1,603 genes from fungi, and the remaining genes from bacteriophages. Of the studied lakes at the two elevations, there were no significantly differences in overall gene richness between the lakes at low and high elevations (*p* > 0.05, Fig. 1a). In all of the detected gene categories (e.g., carbon, nitrogen, phosphorus, and sulfur cycling, organic remediation, metal homeostasis, secondary metabolism, stress response, and virulence), only the richness of stress response genes were found to be significantly higher in the HELs than those in the LELs (*p <* 0.05 in all cases, Fig. 1a), and no significant differences in richness were observed in the rest of the detected gene categories (*p* > 0.05, Fig. 1a).

**Figure 1.**
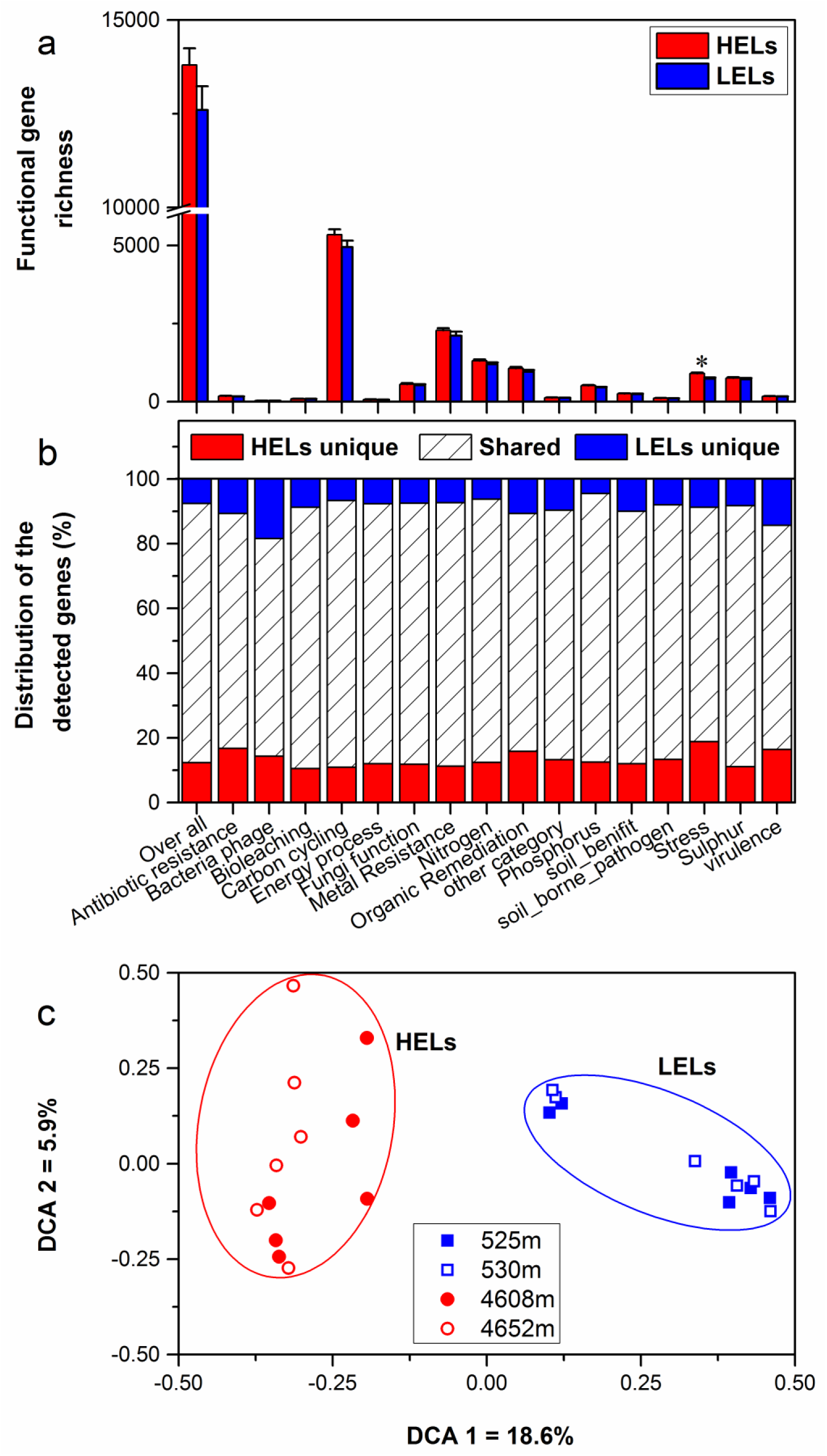
Microbial functional gene richness (a), distribution (shared or unique) of the detected genes (b) and DCA (c) of the microbial functional community structure in the two HELs and two LELs involved in certain biogeochemical cycling processes. All data in panel a and b are presented as the mean ± s.e. calculated from 12 biological replicates. Asterisks (*) above the bars indicate significant (*p* < 0.05) differences. The values on axes 1 and 2 in panel c are percentages of total variations that can be attributed to the corresponding axis.

Microbial functional gene overlap between elevations was also calculated. Only 75.58-87.32% of the genes were shared between the HELs and LELs (Fig. 1b, Table S1). In the LELs, there were in total 7% unique genes, and most of the overall genes (81%) and the genes detected in each gene category (73%-82%) could be found in the HEL (Fig. 1b, Table S1). However, in addition to the genes shared with the LELs, the HELs had another 12% unique genes, most of which were found in stress genes (Fig. 1b).

### Microbial functional gene structure

The MFS differed markedly between the LELs and HELs. The detrended correspondence analysis (DCA) of GeoChip data indicated that the MFSs in the two LELs aligned along DCA axis 1 and those in the two HELs aligned along DCA axis 2 (Fig. 1c). Consistently, three different non-parametric multivariate statistical tests also showed that MFS differed substantially between the HELs and LELs (non-parametric multivariate analysis of variance, ADONIS; analysis of similarity, ANOSIM; multiple response permutation procedure, MRPP; *p <* 0.01 in all cases, Table 1) but not within the two HELs or the two LELs (*p* > 0.05 in all cases, Table 1).

**Table 1.**
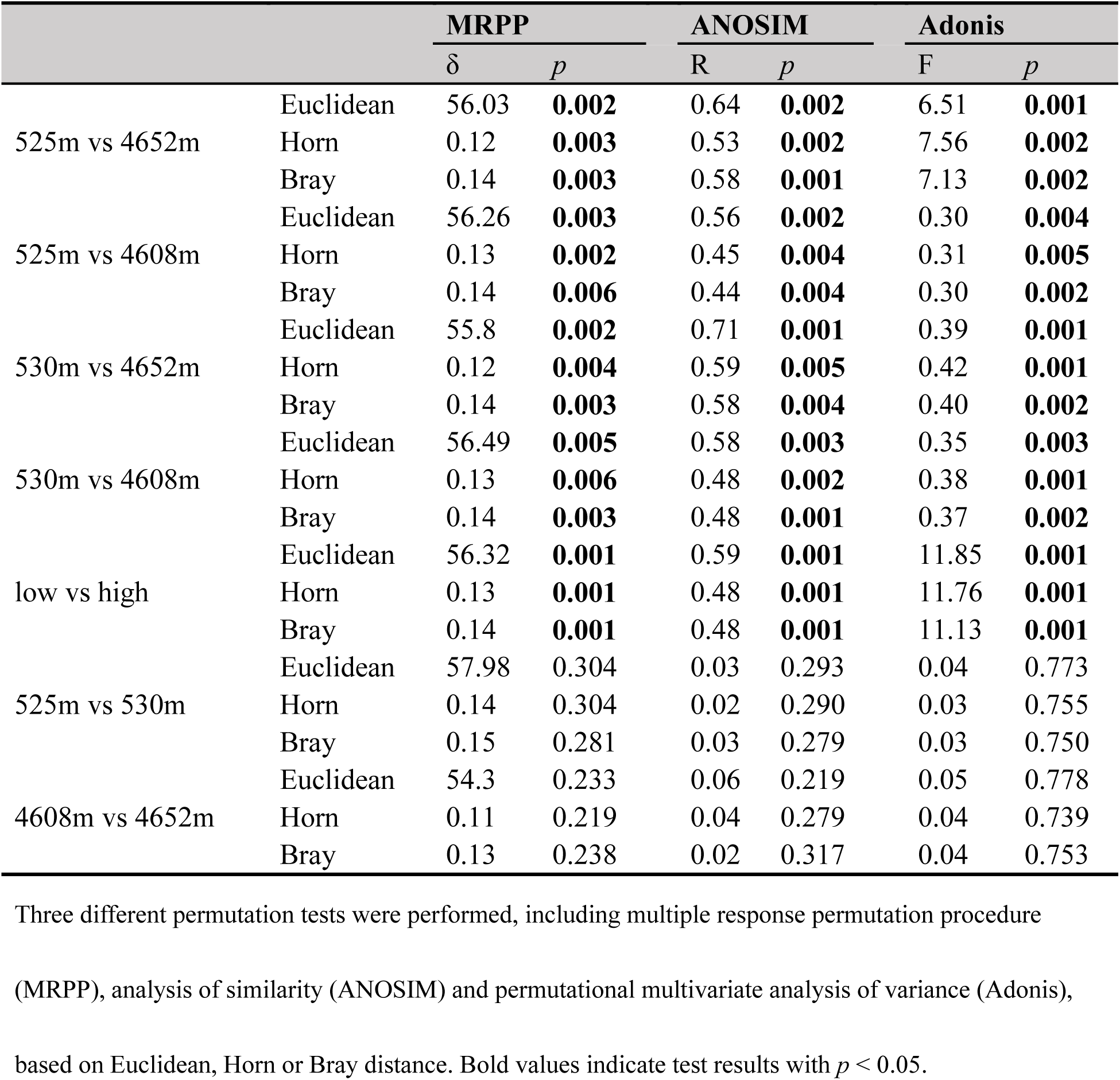
Significance tests of the microbial functional gene structure in lakes at low and high elevations

|  |  | MRPP |  | ANOSIM |  | Adonis |  |
| --- | --- | --- | --- | --- | --- | --- | --- |
| | | $\delta$ | $p$ | R | $p$ | F | $p$ |
| 525m vs 4652m | Euclidean | 56.03 | **0.002** | 0.64 | **0.002** | 6.51 | **0.001** |
|  | Horn | 0.12 | **0.003** | 0.53 | **0.002** | 7.56 | **0.002** |
|  | Bray | 0.14 | **0.003** | 0.58 | **0.001** | 7.13 | **0.002** |
| 525m vs 4608m | Euclidean | 56.26 | **0.003** | 0.56 | **0.002** | 0.30 | **0.004** |
|  | Horn | 0.13 | **0.002** | 0.45 | **0.004** | 0.31 | **0.005** |
|  | Bray | 0.14 | **0.006** | 0.44 | **0.004** | 0.30 | **0.002** |
| 530m vs 4652m | Euclidean | 55.8 | **0.002** | 0.71 | **0.001** | 0.39 | **0.001** |
|  | Horn | 0.12 | **0.004** | 0.59 | **0.005** | 0.42 | **0.001** |
|  | Bray | 0.14 | **0.003** | 0.58 | **0.004** | 0.40 | **0.002** |
| 530m vs 4608m | Euclidean | 56.49 | **0.005** | 0.58 | **0.003** | 0.35 | **0.003** |
|  | Horn | 0.13 | **0.006** | 0.48 | **0.002** | 0.38 | **0.001** |
|  | Bray | 0.14 | **0.003** | 0.48 | **0.001** | 0.37 | **0.002** |
| low vs high | Euclidean | 56.32 | **0.001** | 0.59 | **0.001** | 11.85 | **0.001** |
|  | Horn | 0.13 | **0.001** | 0.48 | **0.001** | 11.76 | **0.001** |
|  | Bray | 0.14 | **0.001** | 0.48 | **0.001** | 11.13 | **0.001** |
| 525m vs 530m | Euclidean | 57.98 | 0.304 | 0.03 | 0.293 | 0.04 | 0.773 |
|  | Horn | 0.14 | 0.304 | 0.02 | 0.290 | 0.03 | 0.755 |
|  | Bray | 0.15 | 0.281 | 0.03 | 0.279 | 0.03 | 0.750 |
| 4608m vs 4652m | Euclidean | 54.3 | 0.233 | 0.06 | 0.219 | 0.05 | 0.778 |
|  | Horn | 0.11 | 0.219 | 0.04 | 0.279 | 0.04 | 0.739 |
|  | Bray | 0.13 | 0.238 | 0.02 | 0.317 | 0.04 | 0.753 |

Three different permutation tests were performed, including multiple response permutation procedure
(MRPP), analysis of similarity (ANOSIM) and permutational multivariate analysis of variance (Adonis),
based on Euclidean, Horn or Bray distance. Bold values indicate test results with $p < 0.05$ .

### Functional genes involved in stress response

Among all functional pathways evaluated, stress response genes were the most variable, most (30 out of 41) of which demonstrated significantly higher normalized signal intensities in the HELs than those in the LELs (*p <* 0.05, Fig. 2). These functional genes that had higher normalized signal intensities in HELs represented all of the detected stress gene categories including pathway-specific genes for response to nutrient and oxygen stress, radiation, cold-shock, heat-shock, and sigma factors, suggesting that the stress response potential in the HELs was greater than that in the ELEs. For example, the normalized signal intensities of the functional genes related to glucose limitation (*bglP*), oxygen limitation (*cydA* and *ahpF*) and osmotic stress (*proW*) increased more than 50% in the HELs comparing to those in the LELs (*p <* 0.05 in all cases, Fig. 2). Similarly, more than 20% of increased normalized signal intensities were detected in the pathway-specific genes related to radiation (*obeE*), heat shock (*grpE, hrcA*, and *dnaK*), nitrogen limitation (*glnR* and *glnA*), phosphate limitation (*pstA, pstB, phoB*, and *pstC*), sigma factors (sigma_32, sigma_24, and sigma_70), and protein stress (*clpC*) in the HELs (*p <* 0.05 in all cases, Fig. 2).

**Figure 2.**
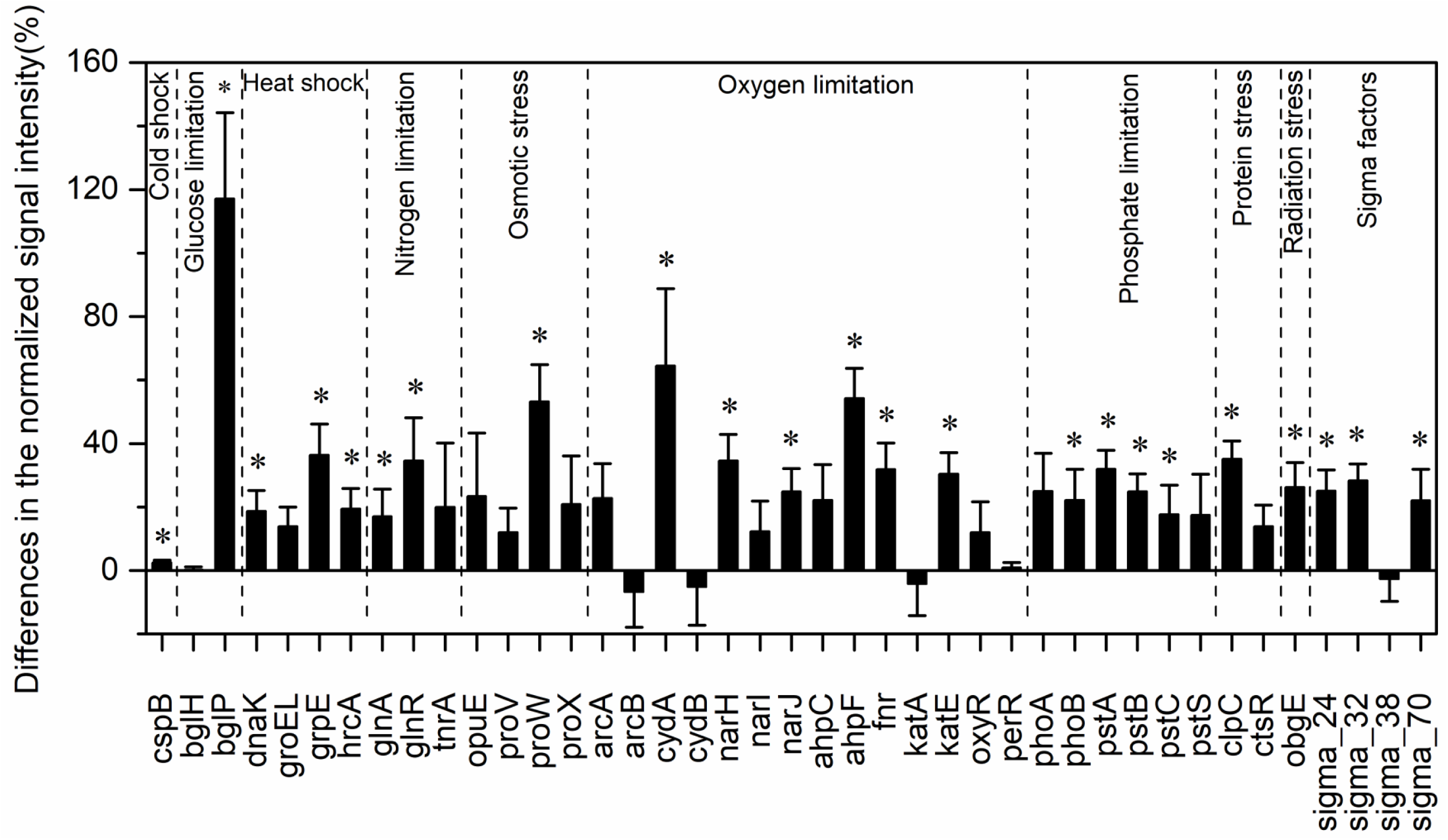
Differences in the normalized intensities of the stress response gene categories between the two LELs and two HELs. All data are presented as mean ± s.e. calculated from biological 12 replicates. Asterisks (*) above the bars indicated significant (*p* < 0.05) differences.

The taxa-function relationships revealed that most of differences in the glucose limitation between the HELs and LELs were exclusively derived from *Firmicutes*, which was not found in the LELs (*p <* 0.05, Fig. S1a). Differences in oxygen limitation between the HELs and LELs were mostly related to *Firmicutes, Betaproteobacteria, Deltaproteobacteria, Gammaproteobacteria*, and Fungi (Fig. S1b). Variations of the pathways related to osmotic stress and nitrogen limitation were both largely attributed to more abundant *Gammaproteobacteria* in the HELs (Fig. S1c and S1e). Differences in heat shock were mostly related to *Actinobacteria* and *Firmicutes* (Fig. S1d, *p <* 0.05 in both cases). Similarly, cold shock that are derived from *Actinobacteria* had higher normalized signal intensities in the HELs than those in the LELs (Fig. S1a, *p <* 0.05). Genes indicative of pathways that respond to phosphate limitation were relatively widespread, but particularly abundant among the *Gammaproteobacteria, Cyanobacteria*, and *Betaproteobacteria* (Fig. S1f). Differences in sigma factors were mostly attributed to *Actinobacteria, Alphaproteobacteria, Cyanobacteria, Deinococcus-Thermus* (primarily *Deinococcus deserti* and *D. geothermalis*), and *Gammaproteobacteria* (Fig. S1g). Differences in protein stress were mostly attributed to *Firmicutes* (primarily *Exiguobacterium sp*., some strains of which can grow within a wide range of pH values, tolerate high levels of UV radiation) and *Verrucomicrobia*, all of which were only detected in HELs (Fig. S1h). Differences in radiation stress were related to the most diverse range of taxa, including *Chlorobi, Cyanobacteria, Deltaproteobacteria, Deinococcus-Thermus, Firmicutes, Gammaproteobacteria*, and *Thermotogae* (Fig. S1i).

### Functional genes involved in carbon cycle

For autotrophic carbon fixation genes, although ATP citrate lyase (*aclB*), carbon-monoxide dehydrogenase (*CODH*), ribulose-1,5-bisphosphate carboxylase/oxygenase (*rubisco*) and acetyl-CoA carboxylase biotin carboxylase subunit (*pcc*) gene groups were observed in both HELs and LELs, the normalized signal intensities of all of them were not significantly different (Fig. 3a, *p* > 0.05 in all cases). Similarly, the metabolic potential of acetogenesis (i.e., formyltetrahydrofolate synthetase (*FTHFS*)) in the Wood-Ljungdahl pathway was also not notably changed with the increase of elevation (Fig. 3a, *p* > 0.05). The same pattern was also detected in methane oxidation genes of *mmoX* and *pmoA* (Fig. 3a, *p* > 0.05 in both cases).

**Figure 3.**
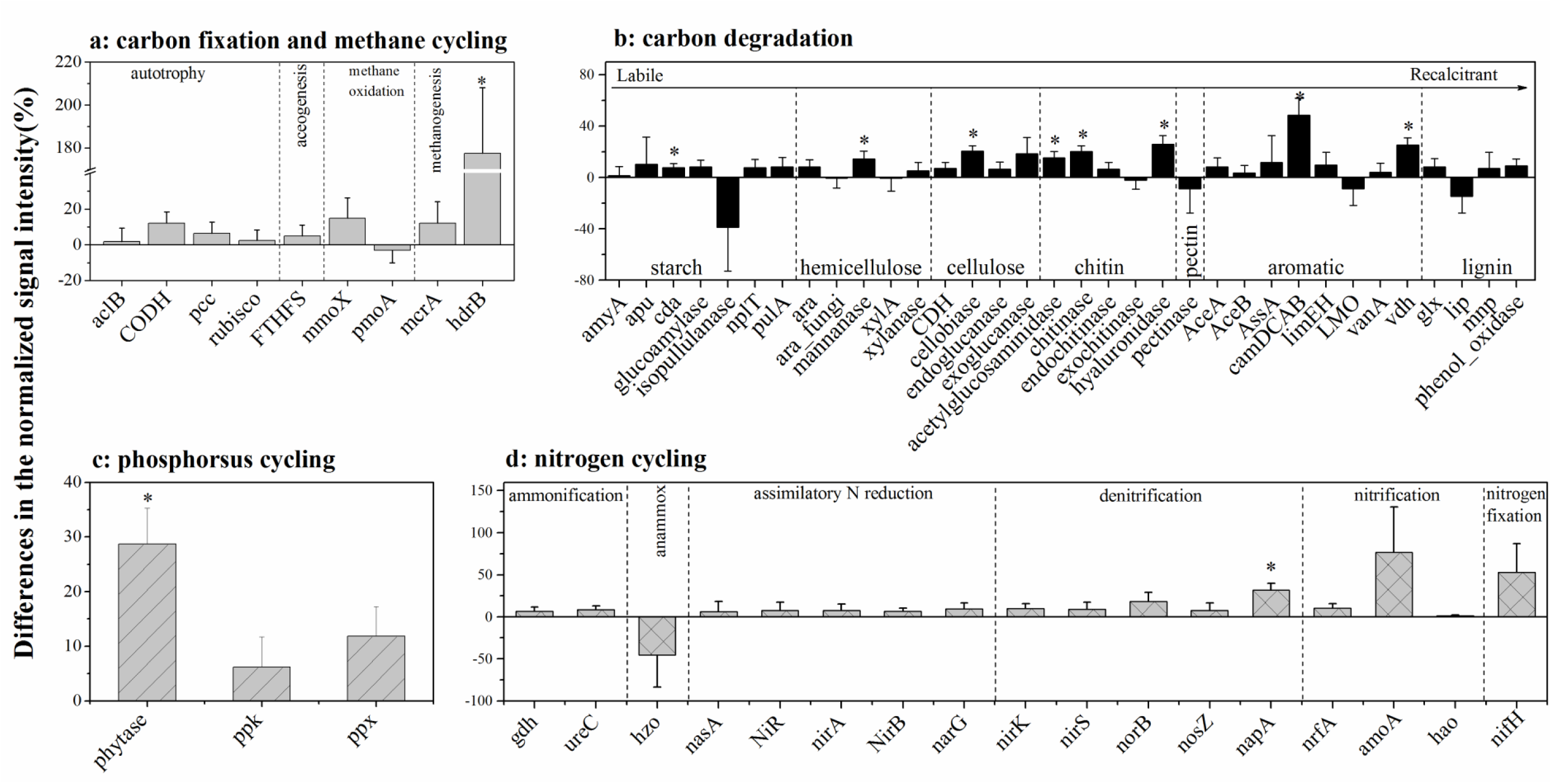
Differences in the normalized intensities of certain biogeochemical cycling processes between the two LELs and two HELs. (a) Each subcategory of the carbon fixation and methane clcling; (b) each subcategory of carbon degradation; (c) each subcategory of phosphorus cycling and (d) each subcategory of nitrogen cycling. All data are presented as mean ± s.e. calculated from biological 12 replicates. * Above the bars indicated significant (*p* < 0.05) differences.

However, the normalized signal intensity of a methanogenesis gene, *hdrB* (methyl-coenzyme M reductase), was nearly two times higher in the HELs than that in the LELs (Fig. 3a, *p <* 0.05). The taxa-function relationships revealed that most of these differences were derived from *Euryarchaeota* and *Crenarchaeota*, and these *Euryarchaeota* (primarily *Archaeoglobus fulgidus* and *Methanolinea tarda*) and *Crenarchaeota* (*Metallosphaera yellowstonensis*) were not detected in the LELs (Fig. S2a).

In contrary to carbon fixing genes, there were eight of the carbon degradation genes, especially the metabolic potentials of recalcitrant carbon such as chitin, and aromatic, showing significantly different normalized signal intensities between the HELs and the LELs (Fig. 3b). Compared with the LELs, the most significantly enhanced carbon degradation in the HELs was related to aromatic metabolizing (*camDCAB*, increased 48%), the second of which included another one aromatic metabolizing (vanillin dehydrogenase (*vdh*), increased 25%), three chitin metabolizing (hyaluronidase, acetylglucosaminidase, and chitinase, increased 26%, 20%, and 15%, respectively), and one cellulose metabolizing (cellobiase, increased 20%). Other more markedly increased normalized signal intensities were one starch degradation gene (cyclomaltodextrinase (*cda*), increased 8%) and one hemicellulose degradation gene (mannanase, increased 14%) (*p <* 0.05 in all cases). These differences were ascribed to some common carbon degradation microorganisms, such as *Firmicutes, Actinobacteria, Gammaproteobacteria, Alphaproteobacteria*, and *Betaproteobacteria* (Fig. S2b-i).

### Functional genes involved in the phosphorus and nitrogen metabolisms

Only the functional potential of *phytase* for phytate degradation was significantly higher in HELs than in LELs among the three P cycling genes (*p <* 0.05, Fig. 3c). This finding was in accordance with higher relative abundance of polyphosphate degradation microorganism from *Gammaproteobacteria, Firmicutes, Alphaproteobacteria, Betaproteobacteria*, and *Bacteroidetes* in HELs (*p <* 0.05, Fig. S2k). Regarding the 17 nitrogen cycling gene groups, there was only one nitrate reduction gene (*napA*) showed remarkably higher normalized signal intensity in the HELs than in the LELs (*p <* 0.05, Fig. 3d). The taxa-function relationship analysis revealed that between HELs and LELs, the notable difference in normalized signal intensity for nitrification was observed primarily in *Actinobacteria* and *Deltaaproteobacteria* (*p <* 0.05, Fig. S2j).

### Microbial functional gene network analyses

In the two networks, we obtained 1,873 nodes and 7,550 links (47% negative and 53% positive) for the HELs and 1,355 nodes and 4,548 links (45% negative and 55% positive) for the LELs (Fig. 4). The typical correlation values of the two constructed functional molecular ecological networks (fMENs) were both 0.98 (Table 2), indicating that the connectivity of the two fMENs fitted the power law model well. The indexes including modularity, clustering coefficients and harmonic geodesic distance were significantly different from those of the corresponding random networks for the lakes at both of the two elevations (Table 2), indicating that the fMENs in both lakes were non-random (*p <* 0.001 in all cases, Table 2). Compared with the LELs, the fMENs in the HELs generally had higher connectivity, shorter geodesic distances, higher clustering efficiencies, and more modules (Table 2).

**Table 2.** Major topological properties of the empirical fMENs and their associated random networks in the LELs and HELs

| Elevation<br>of Lakes<br>(m) | No. of<br>original<br>genes<sup>a</sup> | Empirical networks |  |  |  |  |  |  |  |  | Random networks<sup>c</sup> |  |  |
| --- | --- | --- | --- | --- | --- | --- | --- | --- | --- | --- | --- | --- | --- |
|  |  | Similarity<br>threshold<br>(St) | Network<br>size (n)<sup>b</sup> | Links | R<sup>2</sup> of<br>power<br>law | Average<br>connectivity<br>(avgK) | Harmonic<br>geodesic<br>distance<br>(HD) | Average<br>clustering<br>coefficient<br>(avgCC) | Modularity<br>(No. of<br>modules) | Transitivity<br>(Trans) | Harmonic<br>geodesic<br>distance<br>(HD ±<br>SD) | Average<br>clustering<br>coefficient<br>(avgCC ±<br>SD) | Average<br>modularity<br>(M ± SD) |
| LELs | 1808 | 0.98 | 1355 | 4548 | 0.881 | 6.713 | 6.668<sup>d</sup> | 0.350<sup>d</sup> | 0.790<sup>d</sup>(133) | 0.333 | 3.408 +/-<br>0.014 | 0.029 +/-<br>0.003 | 0.333 +/-<br>0.003 |
| HELs | 2397 | 0.98 | 1873 | 7550 | 0.877 | 8.062 | 6.269<sup>d</sup> | 0.366<sup>d</sup> | 0.865<sup>d</sup>(156) | 0.340 | 3.379 +/-<br>0.009 | 0.023 +/-<br>0.002 | 0.340 +/-<br>0.002 |

2 a. The number of genes used for network construction. b. The number of nodes in a network in a functional molecular ecological network. c. The parameters were 3 generated from 100 times of random rewired networks. d. Significant difference between the empirical network and the random network ( $p < 0.001$ , Z-test).

**Figure 4.**
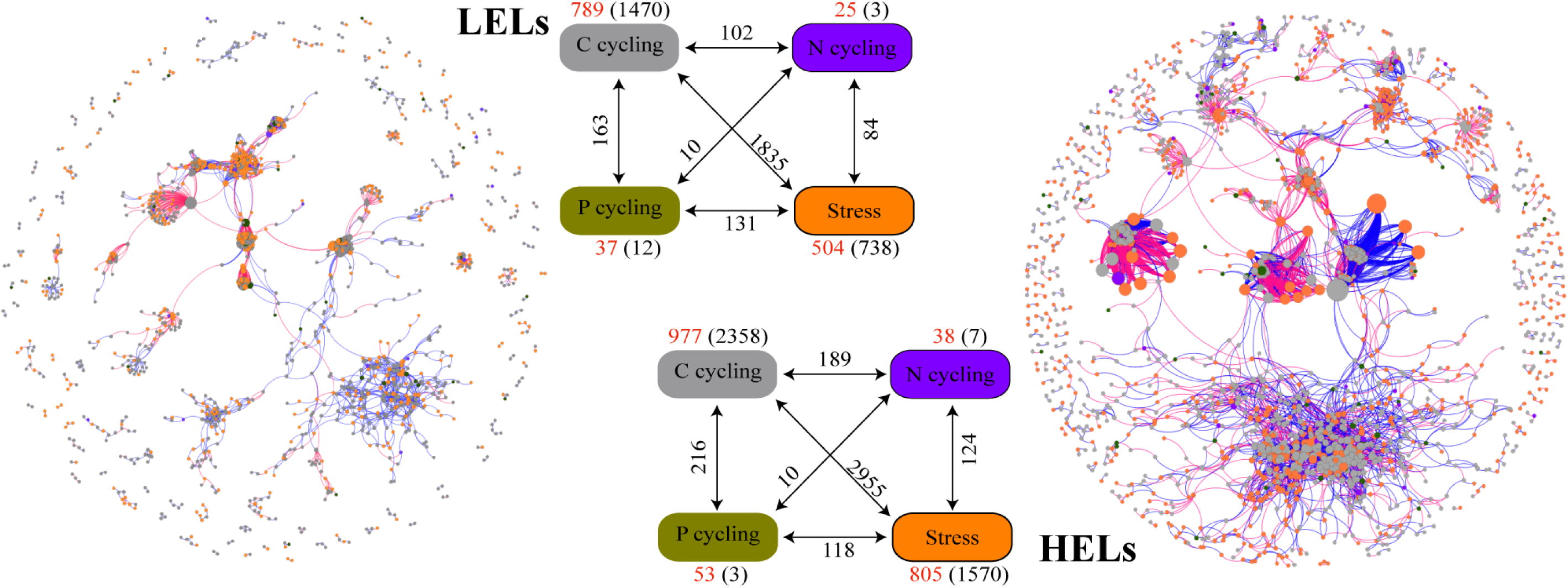
Differences in network interactions of these genes between the two LELs and two HELs. All data in panel a and b are presented as the mean ± s.e. calculated from 12 biological replicates. Network analysis showing the associations within each gene category and associations between different gene categories. A connection indicates a strong (Spearman’s r > 0.8 or r < −0.8) and significant (*p* < 0.01) correlation. A blue line indicates a positive interaction between two nodes, and a red line indicates a negative interaction. The size of each node is proportional to the number of connections (i.e., degree). Colors of the nodes indicate different functional gene categories: orange, stress response; gay, carbon cycling; dark violet, nitrogen cycling; dark goldenrod, phosphorus cycling. Numbers outside and inside parentheses represent the node numbers and inner connections belonging to the corresponding gene category (i.e., there were 504 nodes and 738 inner connections for stress response genes in LELs); the numbers adjacent to edge connections represent cross-gene-category interactions.

More interactions and module hubs in relation to stress response and carbon cycling functional genes were found in HELs network than those in LELs network (Table S3). The hub nodes in the LELs network mainly belong to carbon cycling genes (i.e., carbon degradation subcategory) (Table S3). In contrast, most hub notes and connectors in the HELs network belong to stress response genes (i.e., oxygen stress, oxygen limitation, sigma factors, and phosphate limitation), and some belong to carbon cycling genes (Table S3). No connectors were detected in LELs network while there were two in HELs network, one of which was stress response gene (Table S3).

The top-ranked functional genes in the HELs network were obviously different from those in the LELs (Fig. S3). The top five ranked genes in the HELs belong to stress response genes (*katE, fnr, sigma_24*) and carbon cycling genes (*cellobiase, hyaluronidase*). In the LELs, the top five ranked genes belong to carbon cycling genes (*chitinase, hyaluronidase*), nitrogen reduction gene (*napA*) and stress response gene (*sigma_70*). Moreover, the network interactions of the top five ranked genes in the HELs (Fig. S3a) were much more complex than those in the LELs (Fig. S3b) in terms of network size and connectivity.

### Linkages between MFS and lake environmental parameters

Canonical correspondence analysis (CCA) indicated that four factors including temperature, PO_4_-P, DOC, and turbidity, played key roles in shaping the variance of the MFS variation in the four lakes (for all canonical axes, *p <* 0.05, Fig. 5a). The first two axes explained 19.8% of the observed variation in the composition of the MFS, and 37.6% was explained by the full four canonical axes. All of the four selected factors showed high canonical correlations with the first axis.

**Figure 5.**
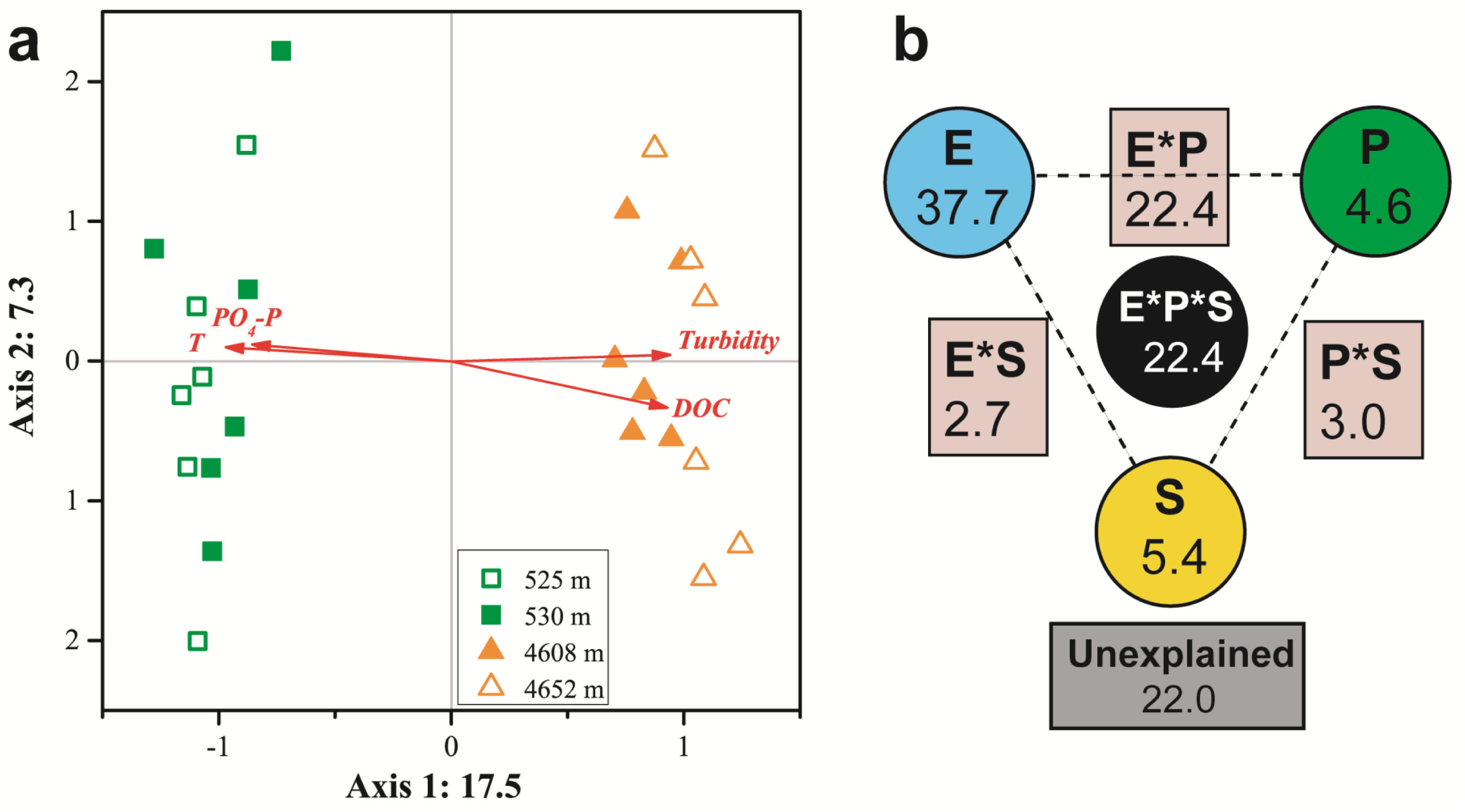
CCA (a) and VPA (b) of relationships between the microbial functional community structure and the environmental variables. P: productivity: Chl *a*, DOC; S: space: elevation and area; E: the rest of the tested environmental variables. Numbers on CCA axis and in VPA diagram show the percentages of explained variations in the microbial functional gene profile. In the VPA diagram, the edges of the triangle presented the variation explained by each factor alone, while the sides of the triangles presented interactions of any two factors and the middle of the triangles represented interactions of all three factors.

The results of variation partitioning analysis (VPA) analysis indicated that environmental heterogeneity was most important in explaining the variance of the MFS variations in the four lakes (Fig. 5b). The three categories of variables, environmental heterogeneity, productivity, and space (elevation and lake area), explained 37.7%, 4.6%, and 5.4% of the total variance, respectively. Interactions among the three groups contributed 30.3% of the total variance. A smaller portion (22%) of the MFS variations could not be explained by the tested environmental variables.

We also revealed that the top 5 modules detected in the HELs and LELs networks showed different relationships with the environmental variables (Fig. S4). In the network of the HELs, modules 1 and 5 were positively correlated with turbidity; modules 3 and 4 were negatively correlated with water temperature, the concentrations of chlorophyll *a* (Chl *a*) and DOC but positively correlated with pH; module 2 was negatively correlated with oxidation reduction potential. However, in the network of the LEL, the largest module 1 showed positive and negative correlations with water temperature and total phosphorus, respectively. Modules 3, and 4, the former of which was negatively correlated with DOC, showed positive correlations with DO. Module 2 did not correlate with any of the investigated environmental variables.

## DISCUSSION

### No differences in overall functional gene richness between the HELs and LELs

In this study, we used the two HELs as a reference to investigate the responses of microbial functional traits to elevational increasing and found that the overall functional gene diversity was not significant higher in the two HELs than in two LELs. This result is different from that found in grass soil where the number of detected microbial genes at the 3,200m site was roughly half of those at the other three higher elevations (3,400 m, 3,600 m and 3,800 m; 20). These differences may mostly be due to covariance of altitude and regional characteristics in terrestrial ecosystems, whereas lakes are more homogeneous (12). For example, pH in soil changed more within the elevational range of 600 m (20) than in lake water within the elevational range of 4,100 m in our study, and pH has been found has an adverse effect on microbial gene diversity (20, 34). Besides this reason, large number of introduced rare bacterial taxa, which were input from lake surrounding catchments or sediments, can increase overall bacterial diversity in small lakes with large catchment area at high evelation (e.g., 12, 35, 36) and are also likely to influence microbial functional gene diversity. Thus, comparing to that in LELs, the microbial gene richness in the two HELs did not decrease but increased a little.

### Enhanced potentials of stress response in the HELs

In contrary to functional gene diversity, microbial metabolic potentials were significant different between the HELs and LELs. The most differences were that the normalized signal intensities of microbial stress genes were higher in the HELs than in LELs (Fig. 1). Previous studies have shown that soil microbial communities at higher elevations had high normalized signal intensities in stress response genes including cold shock, oxygen limitation and osmotic stress genes (20).

Besides theses, in the present study, we also found another 7 subcategories of stress genes, such as nitrogen limitation, phosphate limitation, glucose limitation, radiation stress, heat shock, protein stress, and sigma factors that were higher in normalized signal intensities in the HELs (Fig. 2). These may due to the more significantly changes in environmental factors in the water within a broader elevation gradient and the character of lakes themselves in our present work. With the elevation increased from about 530 m to more than 4,600 m, water temperature dropped nearly 10 times (from 27 °C to 3 °C), Chl *a* decreased more than 20 times, the concentrations of nutrients such as phosphate, nitrate, and ammonium even could be negligible, and DO decreased a half (Table S4). These environmental conditions in the HELs could cause higher microbial metabolic potentials of nitrogen limitation, phosphate limitation, glucose limitation and protein stress genes.

Interestingly, we also detected that the normalized signal intensities of microbial radiation stress genes were significantly higher in the HELs compared with those in the LELs, which were different from the results found in the grass soil in which radiation stress genes were more abundant at the lower elevation (20). This may be due to the fact that the UV radiation is more intensive in surface water than that in the grass soil, where aboveground vegetation prevent UV penetration (20). Our finding of more functional genes with high normalized signal intensities in diverse subcategories of the stress response pathways in the HELs than in the grass soil confirms that recruitment to exposed niches is strongly on the basis of selection for stress tolerance traits and that these microbial communities have a more diverse abilities with which to tolerate harsh environmental conditions than more sheltered niches (22).

Furthermore, higher normalized signal intensities in the microbial functional genes of heat shock, and sigma factors were observed in the HELs than in the LELs, which were also found on the Antarctica rock surface (22) and in the Tibetan mountainous grass soil (20). Mounting evidence suggests that besides protecting microbial cells form environmental insults of sudden temperature increase, the transcriptional activities of heat shock genes decreases but keeps at a steady-state level that is frequently greater than the initial basal level for purpose of assisting microbial growth under non-optimal environmental conditions (37, 38). Evenly, some heat-shock proteins are also required and are abundant during normal growth condition. For example, *GroEL* and *dnaK* genes, play an important role in protein folding even during non-stressed growth conditions, although their action becomes more important during stress (37, 38). Actually, as well as temperature variations, several other stress conditions, such as osmotic changes, desiccation, antibiotics, solvents and heavy metals, are able to elicit heat-shock response (38). In our studied HELs, where are in ice during most time of every year, there are much diurnal temperature variations in summer time, which are likely to cause increase of heat shock genes. Consequently, sigma factors, alternatives of which are the positive transcriptional regulations of shock genes (38), were also increased in the HELs in the present study.

### Enhanced potentials of recalcitrant carbon degradation in the HELs

Though at functional gene category level, there was no significant change in the normalized signal intensities of carbon, phosphorus and nitrogen metabolisms in the HELs (Fig. 1a), at single functional gene level, there were a few of remarkable increase in the normalized signal intensities of carbon degradation genes, especially in recalcitrant carbon degradation genes (Fig. 3b). Differences in the composition of DOC may be a principal factor causing changes in microbial taxonomic structure in aquatic ecosystems (39, 40), which may influence microbial functional metabolic potentials (19). Generally, there are two main carbon sources in lake ecosystems: the terrestrial (allochthonous) inputs of carbon received from landscape around lakes and autochthonous carbon provided by in-lake primary production (41). The relative contribution of allochthonous carbon inputs increase with increasing elevation whereas their liable decrease with increasing elevation (21). In our four studied lakes, the DOC to Chl *a* ratios, which acts as an allochthony indicator (42-44), in the two HELs were averagely 4 times higher than those in the two LELs (Table S4). These low concentration but more recalcitrant DOC in the HELs may promote the microbial functional potentials of recalcitrant carbon degradation, such as, aromatic degradation genes of *camDCAB* and *vdh*, and chitin degradation genes of *acetylglucosaminidase, chitinaseb* and *hyaluronidase* in LELs (Fig. 3b).

### Enhanced interaction among the co-occurring functional genes in the HELs

The functional gene network in the HELs contained more interactions (links) and greater network size, connectivity, and clustering coefficients than those in the LELs, indicating reinforced complexity in the HELs network. Previous studies found that, at continental scale, soil microbial networks detected in the northern China were more complex than those found in the southern China, and demonstrated the impacts of temperature on the complexity of microbial networks at a continental scale (46, 47). Our network analyses indicated that, with the increase of elevation from about 530 m to 4,600 m, the co-occurring functional genes in the microbial functional communities became much more connected and clustered (Table 2, Fig 4), and that temperature and DOC were two of the few most important environmental factors affecting both the networks in the HELs and LELs (Fig. S4). This finding was consistent with the result detected in the acidic mining lakes, in which harsh conditions such as low pH, high concentrations of sulfate and metals, and limited carbon, nitrogen and phosphorus sources resulted in more complex microbial functional gene networks (34). To respond and adapt to the high elevation and its related factors (e.g., low concentration but more recalcitrant DOC, low temperature and large variation of temperature, and high UV radiation), the microbial communities in the HELs not only enhanced their potentials of recalcitrant carbon degradation (such as *cellobiase_bact_arch, chitinaseb* and *hyaluronidase* genes; Fig. 3b) and stress response (e.g., *katE, fnr*, sigma_24 and sigma_70 genes; Fig. 2), but also increased the interaction among them, which played module-hub roles in the network of HELs (Table S3, Fig S3).

## CONCLUSION

Our results indicate that the elevation increasing of about 4,000 m, which relates to changes in different environmental variables, has a strong influence on the profile of microbial functional potentials in lake water column. The harsh conditions (e.g., nutrient limitation, low temperature and large variation of temperature, high UV radiation and more recalcitrant DOC) at high elevation may significantly promote the microbial metabolic potentials among functional genes involved in stress response and recalcitrant carbon degradation in freshwater lakes. Increased numbers in intensity and module hubs of stress response functional genes coupled with enhanced complexity of gene interactions were observed for microbes exposing to the higher elevation stressors. Overall, our results highlights that the limnetic microbial communities could develop functional strategies to cope with the harsh conditions at high elevations.

## METHODS

### Field sampling and measurement of environmental parameters

In this study, we selected two HELs and two LELs at Siguniang Mountain. The Siguniang Mountain, usually referred to as the “Alps of Asia”, is a mountain system located in the eastern region of the Tibetan Plateau in China. Its highest peak has an elevation of 6,250 m, and the upper slopes of the high mountains are mostly covered by snow throughout the year, which offers a desirable elevation gradient as well as a relatively geologically uniform environmental gradient within a small spatial scale (12). The altitudes of two LELs were 525 m and 530 m above sea level, respectively, whereas those of the two HELs were 4,652 m and 4,608 m above sea level, respectively (12). The two LELs (Maojiakou and Baigongyan; Table S4 and Fig. S5) had much higher water temperature, and the concentrations of DOC, Chl *a*, and nutrients than the two HELs (Heihaizi and Baihaizi; Table S4 and Fig. S5). At each lake, the morphometric variables (elevation, latitude, longitude, surface area, maximum depth) of the lake were measured and six uniformly distributed sites were selected for the sampling. From each sampling point, about 1 L and 9 L of water were collected from surface waters (top 50 cm) for the analysis of microbial functional structure and chemical variables, respectively. The methods used for the sampling and determination of the environmental characteristics were described in the reference (12).

### DNA extraction and GeoChip analysis

The total microbial nucleic acids were extracted and purified as described previously (12). GeoChip 5.0, which included more than 57,000 oligonucleotide probes and covered over 144,000 gene sequences from 393 functional gene families involved in carbon, nitrogen, phosphorus, sulfur, stress response, etc. (48), was used to analyze the obtained DNA samples. For each sample, the DNA (500 ng) was labeled with the fluorescent dye Cy-3 (GE Healthcare, CA, USA) by random priming as described previously (49); then the labeled DNA was purified with a QIAquick purification kit (Qiagen, CA, USA) and dried in a SpeedVac (Thermo Savant, NY, USA). The dried DNA was then resuspended in 42 μL hybridization solution containing 1 × HI-RPM hybridization buffer, 1 × a CGH blocking agent, 0.05 μg·μL^-1^ Cot-1 DNA, 10 pM universal standard, and 10% formamide (final concentrations). After mixing completely, the mixture was denatured at 95°C for 3 min and then kept at 37 °C until hybridization. The hybridization was conducted at 67?°C in Agilent hybridization station (Agilent Technologies Inc., Santa Clara, CA). The hybridized GeoChips were scanned by a NimbleGen MS200 scanner (Roche NimbleGen, Inc., Madison, WI, USA) and the images data were extracted using the Agilent Feature Extraction 11.5 software (Agilent Technologies, Inc., CA, USA).

### Data preprocessing

Raw data were uploaded to the microarray analysis pipeline (http://ieg.ou.edu/microarray/) and analyzed as previously described (20, 50). Briefly, the following steps were carried out: (i) select the good quality spots, which with a signal to noise ratio (SNR) more than 2; (ii) the probes that appeared more than approximately 30% of the samples in one lake (two of the six samples in each lake) were selected for subsequent analysis; (iii) for each sample, the signal intensity of each probe was ln (x+1) transformed and then was normalized by dividing the initial signal intensity of each probe by the mean intensity of positive probes in each sample (22). The obtained microarray data matrix (normalized signal intensity) was considered as ‘species’ abundance for microbial functional gene richness and composition analysis (32); and (iv) the normalized signal intensities of all spots of each sample were transferred into relative signal intensities by dividing the normalized signal intensity of a probe by the total normalized signal intensity of a sample. These relative signal intensities were used for further evaluating the microbial metabolic potentials (34). (v) The difference in normalized signal intensity (or the difference in relative signal intensity) of a specific gene family were calculated as: (S_HELs_/ S_LELs_) -1, where the S_HELs_ and S_LELs_ are the total normalized signal intensity of a specific gene family in the HELs and LELs, respectively.

### Network construction

To reduce the complexity of the data sets, we specifically focused on the commonly detected functional genes involved in the key biogeochemical and ecological processes in lakes along elevation gradients, including C, N, P cycling, and stress responses and these genes investigated in the LELs and HELs were selected for the construction of the fMENs, which were based on RMT-based approach using a publicly Molecular Ecological Network Analysis Pipeline (MENAP) (http://ieg2.ou.edu/MENA/, 32, 51). Briefly, the fMEN was constructed based on the Pearson correlation matrix calculated by the correlation coefficient (r value) between each of the two detected genes (51). Then, a series of thresholds was applied to the matrix and the similarity threshold (st) were kept or calculating matrix eigenvalues using RMT-based approach (52). During this calculating, the most suitable st was selected to obtain the Poisson distribution of the calculated eigenvalues, which indicates the nonrandom properties of a complex system (52). Each network was then separated into modules by fast greedy modularity optimization. For each network, 100 corresponding random networks were then generated with the same network size and an average number of links. The differences in the indices between the constructed and random networks were determined by the statistical Z test. Meanwhile, the Student *t* test was employed using the standard deviations derived from corresponding random networks to compare the network indexes in the lakes at different elevations (32).

The Cytoscape 3.6.1 software (53) and Gephi 0.9.2-beta (54) were applied to visualize the network of the nodes.

### Identification of node roles

The topological roles of each functional gene can be decided by its within-module connectivity (Zi) and among-module connectivity (Pi) (55). The nodes in a network could be organized into the following four categories: network hubs (Zi > 2.5 and Pi > 0.62), connectors (Pi > 0.62), module hubs (Zi > 2.5) and peripherals (Zi < 2.5 and Pi < 0.62), the former two of which have important roles in network topology (32).

### Statistical analyses

Using the microarray analysis pipeline (http://ieg.ou.edu/microarray/), Student’s t-test was carried out to determine the differences in the functional gene diversity and the relative signal intensity of each functional gene category and certain subcategories or phylogenetic groups in the two types of lakes. Three non-parametric multivariate statistical tests, including ADONIS (‘Adonis’ function), ANOSIM (‘anosim’ function) and MRPP (‘mrpp’ function), and DCA were performed to determine the differences in the microbial functional gene structure. All of these four analyses were conducted using the vegan package (version 2.3-0, 56) in the R statistical computing environment.

CCA was performed to detect the relationships between MFS and environmental parameters because the length of the first DCA axis run on the species data was >2. Forward selection was carried out to determinate the significant factors in the CCA model by 999 simulated permutations (*p* < 0.05). All the environmental variables were log (x+1)-transformed except for the values of water temperature and pH. In addition, VPA were also performed to identify individual and interactive contributions of productivity (Chl *a* and DOC), space (elevation and lake area), and the rest of the tested environmental heterogeneity to MFS (12). CCA and VPA were performed using the vegan package in R.

The correlation coefficients between modules and environmental factors were tested on MENAP. The module eigengene E (the first principal component of modules) (57) of the top modules for both the HELs and LELs networks were calculated, and then their relationships with environmental variables were estimated using Spearman’s rank correlation test.

## ACKNOWLEDGEMENTS

We thank Joe Zhou and Ruizhu Huang for their support on Geochip and data analysis of the samples. This work was supported by the Chinese Academy of Sciences (QYZDJ-SSW-DQC030), National Science Foundation of China (91951000, 41621002) and the Second Tibetan Plateau Scientific Expedition and Research Program (2019QZKK0503).

